# Women in neotropical science: Gender parity in the 21^st^ century and prospects for a post-war Colombia

**DOI:** 10.1101/477422

**Authors:** Camilo López-Aguirre

## Abstract

An increasing amount of research has focused on studying the drivers shaping demographics in science. As a result, we now have a better idea of the current state of gender disparity in science, which remains widespread worldwide. However, fewer studies and limited data have restricted our understanding of this phenomenon in the Neotropics, a highly important region in terms of cultural and biological diversity. Despite a civil war that lasted more than five decades and produced eight million victims (half of them women), Colombia is the fifth country with the highest scientific production in Latin America and the Caribbean, as well as the second most biodiverse country in the world. In order to evaluate the status of gender parity in science in Colombia throughout the 21st century, data of science demographics was gathered covering the 2000-2017 time period. Percentage of women in science was decomposed by research area, researcher rank level and education level. Gender disparity was also estimated for changes in average age, access to scholarships for postgraduate studies, and number of doctoral graduates. Finally, using logistic function modelling, temporal projections into the future were performed, in order to estimate how long could it take to reach gender parity. Of six research fields, medical and health science is the only one to have reached gender parity (55.99%), although it is also the only one showing a steady decrease in women representation across time. On the other hand, engineering, humanities and natural sciences had the lowest percentages of female representation (19.89%, 30.02%, and 30.21%, respectively). Female researchers were on average younger than male researchers, and they also showed a decreasing presence as they move upward to more senior levels, exemplifying the ‘leaky-pipeline phenomenon’ common in science. More men were observed both as scholarship awardees for doctoral studies, and as doctoral graduates, indicating that obtaining a doctoral degree could be a major limiting factor for women in science. Possible drivers of these results are analysed, suggesting that a combination of lack of research funding, insufficient legal framework, pre-existing biases, and poor protection of women’s rights inhibits female participation in science. Based on logistic function modelling it is estimated that, without any action to change current trends, it could take between 10 (humanities) and 175 (engineering) years to reach gender parity across all research areas.

## Introduction

Science has grown in the diversity of fields and approaches in which it operates, to the point of including the study of the scientific endeavour itself. The last two decades have witnessed an increased interest in studying the demographics of the workforce in research and development (R&D) (Kern et al., 2015), raising concerns on topics such as inclusivity (Ceci & Williams, 2011), mental health (Evans et al., 2018), multiculturality (Bernard & Cooperdock, 2018), gender parity (Smeding, 2012), and pay gap (Franco-Orozco & Franco-Orozco, 2018), among others. As a consequence, there are now numerous ongoing debates in an effort to improve the working conditions on all of the different branches of R&D (Stirling, 2007). Historically, science has been traditionally patriarchal, favouring the proliferation of a false assumption that men are innately more well-suited for R&D. In an effort to disprove this idea and encourage higher women representation in R&D, studies have focused on understanding the prevalence of subconscious bias and unfavourable conditions for women in Science, Technology, Engineering, Mathematics and Medicine (STEMM) (Christie et al., 2017; van den Besselaar & Sandstrom, 2017). As a result, for the first time in history we have quantitative data to analyse the current state of the R&D workforce (Ceci et al., 2009; Ovseiko et al., 2016), allowing us to make more informed decisions at an individual, institutional and governmental level. Although major improvements have been accomplished on gender parity in undergraduate education, where women are increasingly studying science-related degrees (Franco-Orozco & Franco-Orozco, 2018; Valentova et al., 2017), women representation at postgraduate studies and research positions steadily decrease as they pursue research-intensive careers at more senior levels (Pell, 1996). Moreover, this phenomenon has shown differential trends depending on the area, with engineering and physics consistently showing more dramatic gender disparities (Holman et al., 2018). Gender disparity in science has also been reported in a myriad of variables other than workforce representation, such as conference participation (Débarre et al., 2018; Jones et al., 2014), editorial boards’ composition (Cho et al., 2014), sentiment towards science communicators (Amarasekara & Grant, 2018), grant success rate (Ley & Hamilton, 2008; Pohlhaus et al., 2011; van der Lee & Ellemers, 2015), and papers’ authorship (Holman et al., 2018), exemplifying the complexity of this issue. Sexual harassment has especially impacted the scientific community as reports have shown a hostile environment for women in research and academia across the globe (National Academies of Sciences, Engineering & Medicine, 2018), with several instances where senior researchers were involved in longstanding cases of sexual misconduct (Wadman, 2018).

According to the UNESCO Institute for Statistics (UIS), only a third of the global workforce in science are women (UNESCO Institute of Statistics, 2018). Myanmar and Bolivia are the countries with the highest percentage of women in science (83% and 63%, respectively), whereas at a regional scale Central Asia and Latin America and The Caribbean are world leaders in gender parity in science with 48% and 45% respectively (UNESCO Institute of Statistics, 2018). Nevertheless, loss of gender parity at postgraduate and more senior levels seem to be also present in these countries and regions. By 2016, a report led by the Interacademy Partnership concluded that on average women represented only 12% of the members of 69 Academies of Science worldwide (The Interacademy Partnership, 2015). Science academies in Latin America and The Caribbean had the highest women representation (17%) followed by North America (15%) and Central and Eastern Europe (13%). Female researchers have also been found to be less internationally mobile, less likely to participate on international research collaborations, and less likely to publish papers as first authors, especially on high impact journals (Elsevier, 2015). Projections on authorship in scientific publishing suggest that it could take more than 100 years to reach gender parity in areas such as statistics and physics (Holman et al., 2018). Widespread gender disparity in science is in conflict with findings indicating that research impact is not gender-related, and that female researchers represent a larger proportion of interdisciplinary research outputs (Elsevier, 2015). Significant efforts to promote gender parity include the creation of institutions such as the L’Oreal-UNESCO For Women in Science program established in 1998, the Athena SWAN accreditation program established in 2005 by the British Equality Challenge Unit, and the Science in Australia Gender Equity (SAGE) established in 2015 as a partnership between the Australian Academy of Science and the Australian Academy of Technology and Engineering.

Despite all these efforts, a major impediment to the implementation of policies that promote gender parity is the lack of information on the extent and magnitude of gender disparity in science at local scales, especially in nations with low R&D expenditure. Only until recently, institutions in countries with low R&D expenditure have started meaningful efforts to establish a baseline understanding of the prevalence of gender disparity (Franco-Orozco & Franco-Orozco, 2018; Valentova et al., 2017), limiting our understanding of the historical trajectories of women participation in science.

Regardless of Latin America’s relatively good performance in global indexes of gender parity in science, Latin American female researchers still face many challenges when pursuing a career in science (Daza & Bustos, 2008; Franco-Orozco & Franco-Orozco, 2018; Valentova et al., 2017). Within the Latin American context, over the last couple of decades most of the scientific output was produced by five countries: Brazil, Mexico, Argentina, Chile and Colombia (Scimago, 2018). According to the Scimago Country Rankings, of all the scientific publications authored by researchers in Latin America from 1996-2017, 88.37% was produced between these five countries (Scimago, 2018). Despite these five countries representing the scientific powerhouse of Latin America, most of them suffer from lack of funding (Brazil being the only one with a R&D expenditure >1% of GDP) (World Bank, 2018), gender disparity (Argentina being the only one that has achieved gender parity) (UNESCO Institute of Statistics, 2018), and gender pay gap (Franco-Orozco & Franco-Orozco, 2018). A recent study evaluating women participation in scientific publishing worldwide over the last decade found that none of these five countries has reached gender parity, estimating that women participation in Argentina and Colombia will decrease, moving away from gender parity (Holman et al., 2018). Colombia’s R&D expenditure has consistently stayed below 0.40% of the GDP (the lowest amongst these countries), with a steady reduction of expenditure in recent years. Science gender parity in Colombia remains elusive with women representing only 38% of researchers, and 14% of the active members of the Colombian Academy of Exact, Physical and Natural Sciences. Gender salary gap in postgraduate graduates in the last decade exceeded 30% in areas such as medical sciences and engineering (Franco-Orozco & Franco-Orozco, 2018). Nevertheless, gender parity in education in Colombia has improved since 2000 both at undergraduate and postgraduate level, where between 2011 to 2014 women represented 55, 47.7 and 38.3% of the graduates at undergraduate, masters and doctoral levels, respectively (Franco-Orozco & Franco-Orozco, 2018).

The signing of a peace deal in 2016 to put an end to the armed conflict between the Colombian government and the Revolutionary Armed Forces of Colombia (FARC) could facilitate the reordering of national priorities (Ocampo-Peñuela & Winton, 2017; Salazar et al., 2018), providing an opportunity to improve the working conditions of women in science (Baptiste et al., 2017). The internal civil war in Colombia, the longest-lasting armed conflict in the western hemisphere, left over 260,000 deaths and 7 million people displaced (Daza & Bustos, 2008; Overseas Development Institute, 2015; Oxfam International, 2017; Unidad de Víctimas, 2017). The severity of the conflict also added additional limitations to scientific efforts in Colombia (Augusto et al., 2017; Canavire-Bacarreza et al., 2018; Sierra et al., 2017). Vast regions of pristine native ecosystems remained inaccessible to researchers for 50 years, risking kidnapping and assassination (Baptiste et al., 2017). The combination of limited funding and bad working conditions also created a collateral brain drain, with Colombia having one of the lowest rates of doctoral graduates in Latin America (8 per million inhabitants) (UNESCO Institute of Statistics, 2018), and a significant number of Colombian doctoral graduates residing overseas (El Tiempo, 2017). Post-conflict efforts to strengthen scientific production in Colombia have shown decisive advances, such as the Colombia BIO programme, a series of scientific expeditions inventorying unexplored ecosystems in Colombia, that have resulted in the description of over 100 species previously unknown to science (COLCIENCIAS, 2016).

The combination of challenges and opportunities that science in Colombia is facing reinforces the need to have a detailed (Ocampo-Peñuela & Winton, 2017), overarching perspective of the current status of gender parity in science, in order to diagnose recent trends and inform future efforts and policies that would focus on promoting the participation of women in R&D. The purpose of this study was to assess the demographics of the scientific workforce in Colombia in the last two decades. To do so, I dissected women participation in science by estimating gender parity across research fields, research rank, and training level from 2000 to 2017. Furthermore, I estimated changes in age distribution across genders and incorporated estimates of gender parity in access to education scholarships and number of postgraduate graduates. Finally, I modelled the trajectory of women participation across time in order to provide a rough prediction of the year when gender parity will be achieved at different levels. Based on the local information available, and global patterns of gender parity in science, it is expected to find greater women underrepresentation in engineering-related research fields, as well as a decrease in women representation at higher education levels and more senior levels of research.

## Material and methods

### Data acquisition

Demographics data of the Colombian workforce in science was retrieved from the UNESCO Institute of Statistics (UNESCO Institute of Statistics, 2018), the Network for Science and Technology Indicators –Ibero-American and Inter-American– (RICYT, 2018), the Colombian Science and Technology Observatory (OCyT; Observatorio Colombiano de Ciencia y Tecnología) (OCyT, 2018), and the Science in Numbers data repository (SN) by the Administrative Department of Science, Technology and Innovation (COLCIENCIAS) (COLCIENCIAS, 2017). SN aggregated individual data for 39,342 researchers from the SCIENTI online platform between 2013 to 2017. SCIENTI was developed by COLCENCIAS as an online registry of CVs of individual researchers and research groups in Colombia. Demographic data was sorted based on research area, training level, researcher rank level, doctoral graduates and access to scholarship grants. Overall percentage of women in science (between 2000 and 2015) and by research field (2006 and 2015) was collected from the UIS and RICYT, respectively. Gender parity was decomposed following the OECD classification of research areas: Agricultural sciences, engineering, humanities, medical and health sciences, natural sciences and social sciences. Data for age distribution, training level and researcher rank level between 2013 and 2017 were retrieved from the SN by COLCIENCIAS. Training level was classified in five different groups: Undergraduate, diploma, masters, PhD and postdoctoral; whereas researcher rank level was classified into four groups: Junior, associate, senior and emeritus. Data on gender parity in doctoral graduates was assessed based on data retrieved for 2006 to 2015 from the OCyT and RICYT. Numbers of granted scholarships by gender from 2006 to 2015 presented in this study were collated by the OCyT from different sources and retrieved from its 2016 Science and Technology Indicators report.

### Statistical analysis

To test whether investment in R&D correlates with women representation in science, a linear model was used, based on data of R&D expenditure (as a percentage of GDP) gathered from the World Bank Open Data (World Bank, 2018). To build a temporal perspective of women in science in Colombia, the representation of women in science was calculated annually for each variable, and represented graphically both as a percentage, and as number of researchers for a given year. In order to test whether access to education associates with the number of female researchers in science, linear regression modelling was used to quantify the correlation between the percentage of female researchers and the percentage of female doctoral graduates across time. Finally, to project how long could it take to reach gender parity, a logistic regression model of proportion of women in science across time was used (assuming a sigmoidal relation between gender ratio and time; see Holman et al. (2018)), predicting the year in which the percentage of women reaches 50%. Using a logistic model allows for a non-linear growth rate that plateaus at a maximum value of 1, indicating in this case the complete loss of one gender (Holman et al., 2018). 95% confidence intervals were calculated based on 1,000 bootstrap iterations. All analyses were performed in R version 3.3.2. Logistic modelling was performed with the glm function, and temporal projections of future proportion of women in science were performed using the predict function of the stats package.

## Results

Overall, women representation in science has increased in Colombia across the 21st century. Over the last 15 years, women representation grew by 4.69%, going from 33.71% by 2000, to 38.40% by 2015 (Fig. 1). Raw number of researchers showed the same tendencies across genders, with three periods of increase in the number of researchers (2000-2003, 2004-2011, and 2014-2015), and two periods were the number of researchers decreased (2003-2004, and 2011-2014).

**Figure 1.**
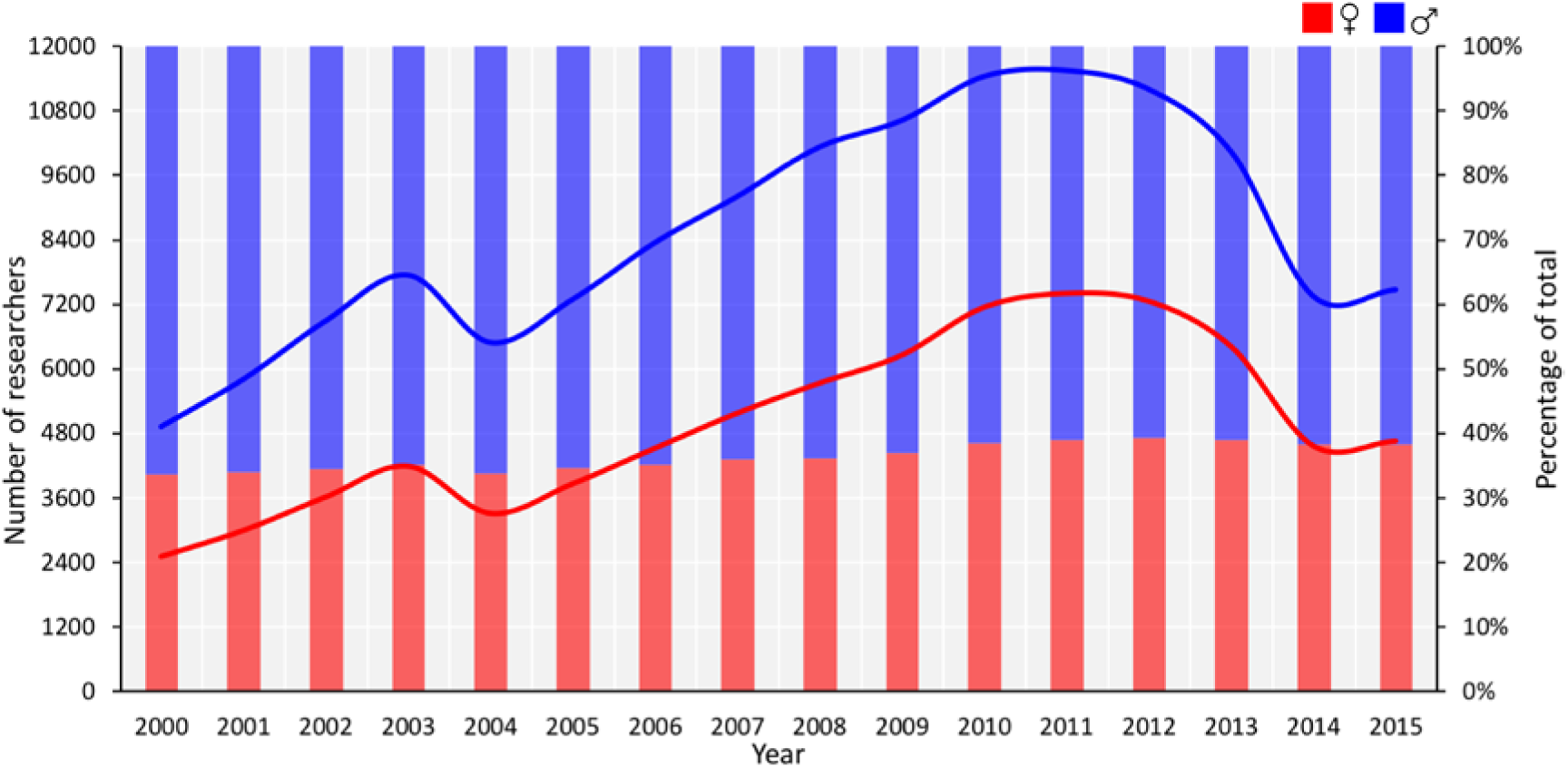
Women (red) and men (blue) participation in science in Colombia between 2000 and 2015, represented as number of researchers (lines) and percentage of total researchers (bars).

Across research fields (Fig. 2), averaging the 2005-2015 period, medical and health sciences showed the highest percentage of female researchers (the only research field that reached gender parity), followed by social and agricultural sciences (55.99%, 44.20%, and 35.91%, respectively). Contrastingly, engineering showed the lowest average of women representation, followed by humanities and natural sciences (19.89%, 30.02%, and 30.21%, respectively). Temporal trends reveal that the humanities had the highest increase of women representation with an increase of 13.85% between 2005 and 2015 (Fig. 2C). Natural sciences and engineering also showed an increase in women representation for the same time period (7.34% and 4.28%, respectively; Fig. 2B and E). Agricultural and social sciences showed almost no change across time (0.22%), reflecting temporal unsteadiness in agricultural sciences where women participation grew initially, and decreased subsequently, and temporal invariability in social sciences, ranging between 43.01 and 46.16% (Fig. 2A and F). Despite having reached gender parity, medical and health science is the only field showing a temporal decrease in women representation, losing 4.94% between 2005 and 2015 (Fig. 2D).

**Figure 2.**
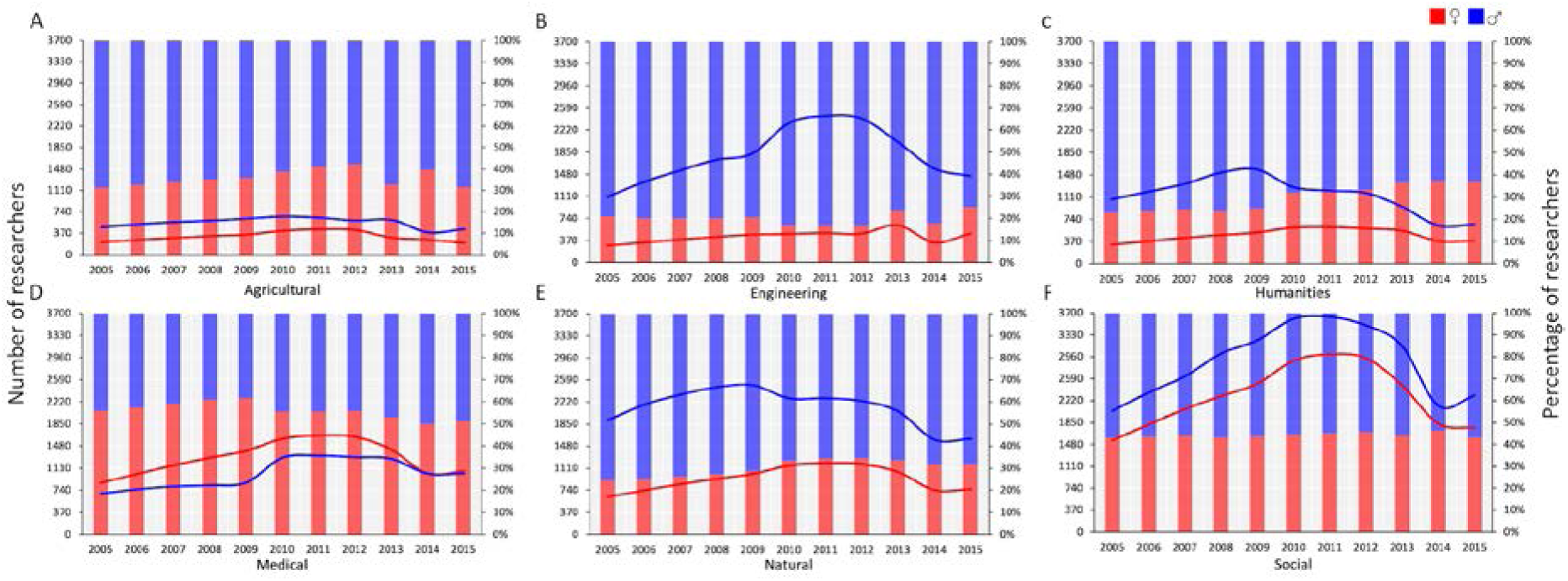
Women (red) and men (blue) participation in science in Colombia between 2000 and 2015, represented as number of researchers (lines) and percentage of total researchers (bars). Results are divided into six different research fields: Agricultural (A), engineering (B), humanities (C), medical (D), natural (E), and social (F).

Between 2013 and 2017, ages 25-45 represented more than half of researchers across genders, with ages 35-40 being the most frequent (Fig. 3). Average age of researchers in Colombia has decreased for both genders between 2013 and 2017, women having the lowest average (44.39 for women and 45.49 for males). Age of female and male researchers decreased by 2.24 (46.06 in 2013 to 43.82 in 2017) and 2.19 years (47.16 in 2013 to 44.97 in 2017), respectively. Lowest age recorded for SCIENTI-registered individuals was lower for males (15-20) than females (20-25). Similarly, Highest age recorded for SCIENTI-registered individuals was higher for males (90-95) than females (80-85).

**Figure 3.**
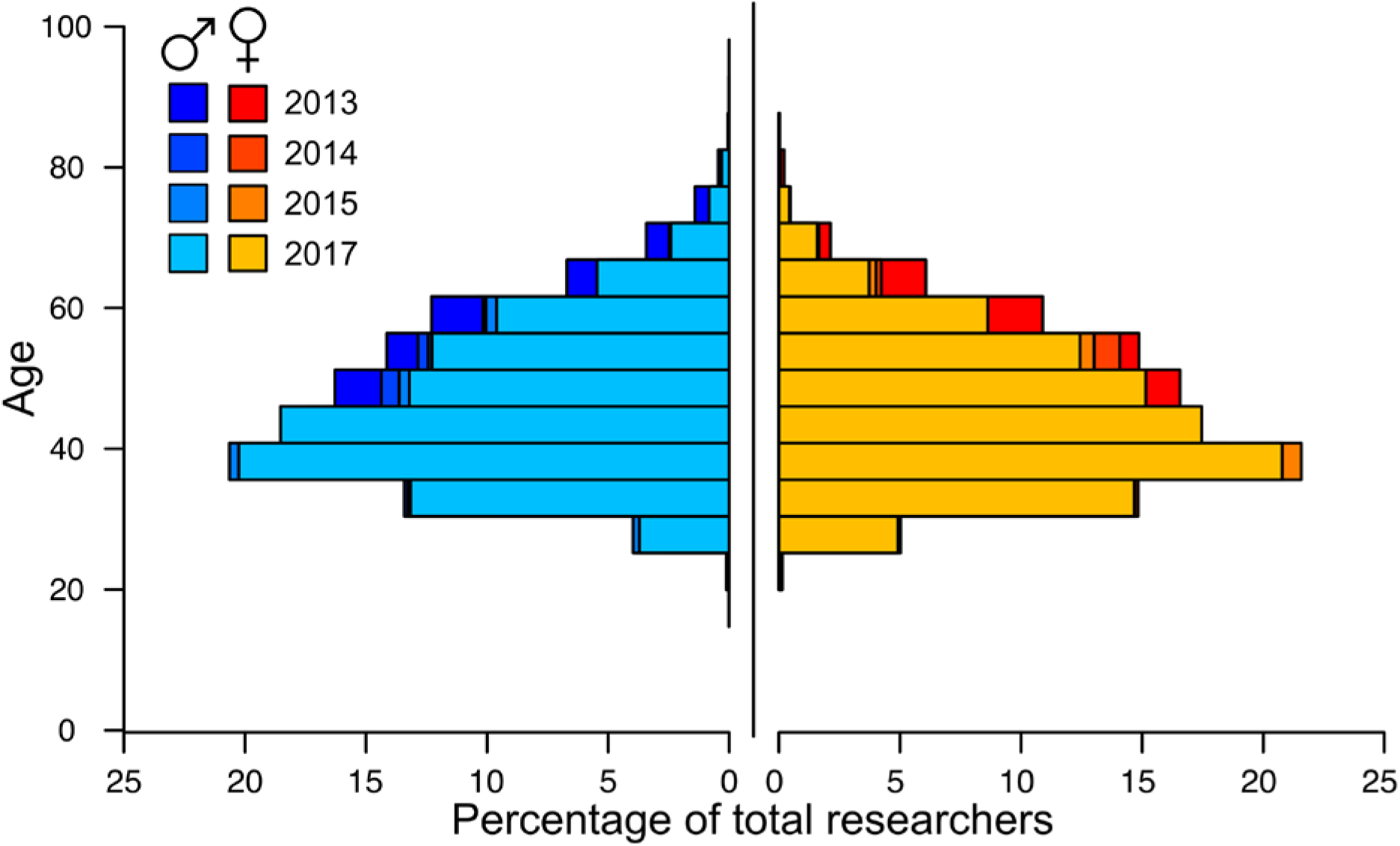
Age distribution of male (left) and female (right) researchers represented as percentage of total researchers of each gender per year, between 2013 and 2017. Shades of red-to-yellow (female) and blue-to-turquoise (blue) represent temporal changes in age distributions.

Proportion of training level of Colombian researchers in the SCIENTI platform showed similar patterns for both genders (Fig. 4). Across time, doctoral degrees were the most abundant, representing more than half of researchers, followed by master’s degrees (20 to 40%), postdoctoral positions (5 to 12%), diplomas and undergraduate (both under 5%). Between 2013 and 2014, percentage of researchers of both genders with a doctoral degree dropped on average by 10%, whereas researchers with master’s degrees increased approximately by the same amount. Furthermore, the proportion of researchers with doctoral degrees was higher for males than for females, and the proportion of researchers with master’s degrees was higher for females than for males. Researchers with postdoctoral-level training was higher for males than females.

**Figure 4.**
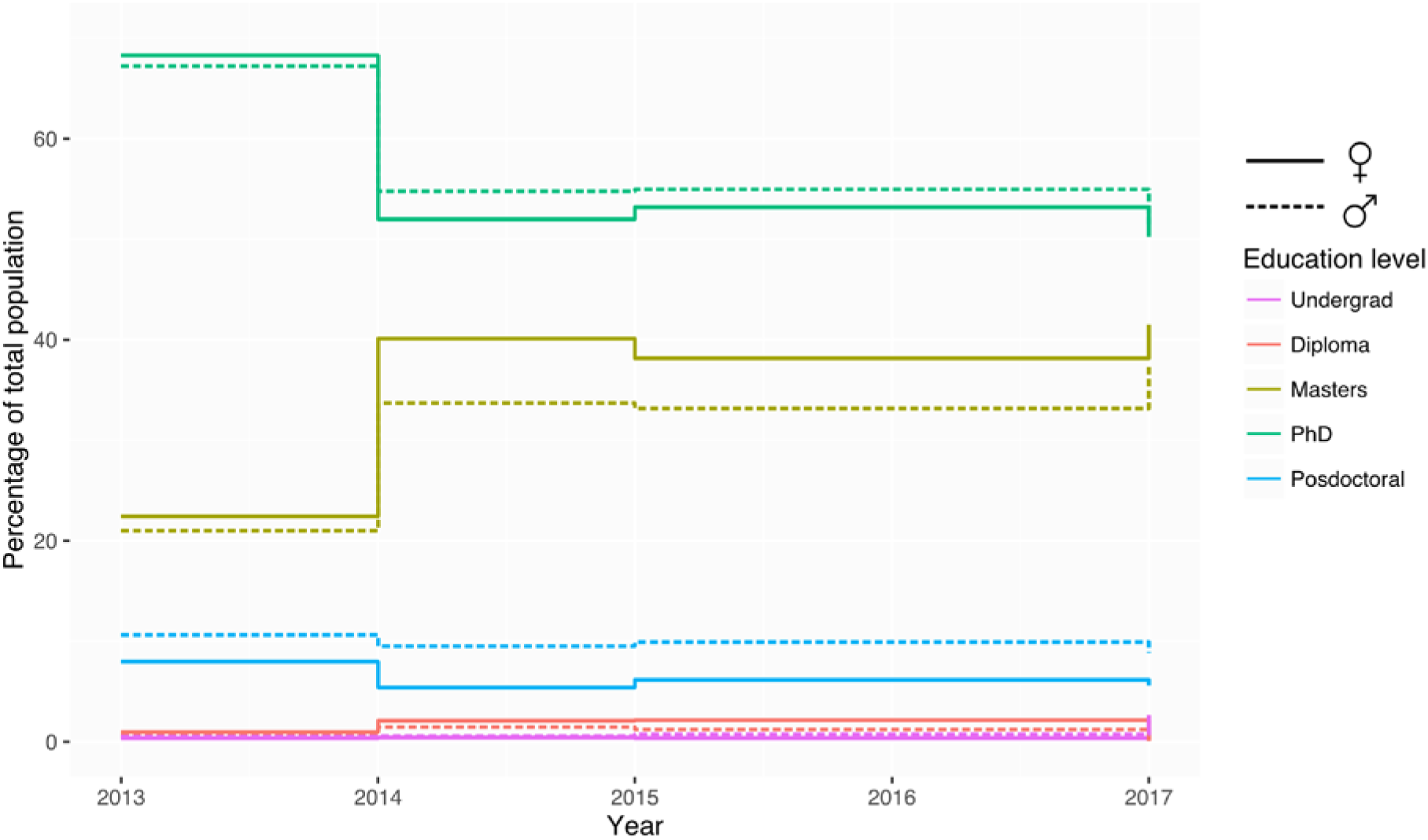
Temporal changes in the proportion of female (solid lines) and male (dashed lines) researchers with different education levels between 2013 and 2017.

Women were underrepresented across researcher rank levels and across time, with a marked widening of the gender gap at more senior research rank level (Fig. 5). Averaging the 2013-2017 period, women represented 37.72%, 35.05%, 25.96%, and 21.51% of junior, associate, senior and emeritus researchers, respectively. Women representation has increased across all rank levels in the time period analysed, with the highest rates of increase in the junior (6.03%) and emeritus (2.49%) levels, followed by the senior (1.13%) and associate (0.98%) researcher rank levels. The only decrease in the number of researchers was evident for male junior researchers between 2013 and 2014 (Fig. 5A). Data for the emeritus rank level was lacking for 2013 and 2014 (Fig. 5D).

**Figure 5.**
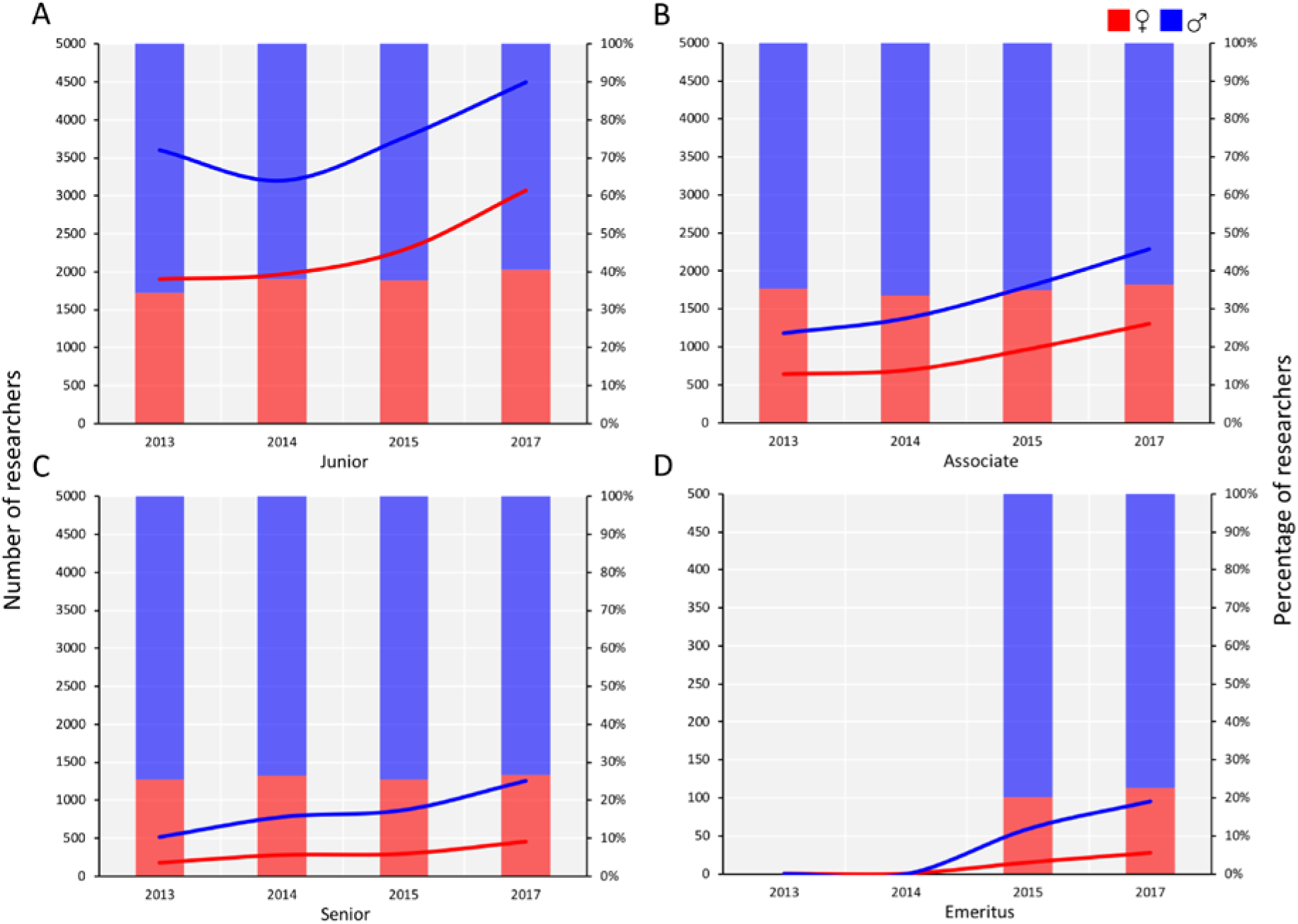
Female (red) and male (blue) representation across researcher rank levels in Colombia between 2000 and 2015, represented as number of researchers (lines) and percentage of total researchers (bars). Results are divided into six different research fields: Junior (A), associate (B), senior (C), and emeritus (D).

Women representation in doctoral graduates in Colombia between 2006 and 2015 remained below parity across research areas (Fig. 6). Medical sciences had the highest average percentage of female doctoral graduates (48.65%), next to social (43.20%), agricultural (36.79%), natural (35.98%), humanities (35.03%), and engineering sciences (25.71%). The highest increase in the proportion of female doctoral graduates was found in the medical (27.77%, Fig. 6D), agricultural (16.92%, Fig. 6A), and social sciences (12.68%, Fig. 6F). Contrarily, humanities (2.95%), engineering (3.29%), and natural sciences (6.86%) showed the lowest increase in the proportion of female doctoral graduates (Fig. 6B-C,E).

**Figure 6.**
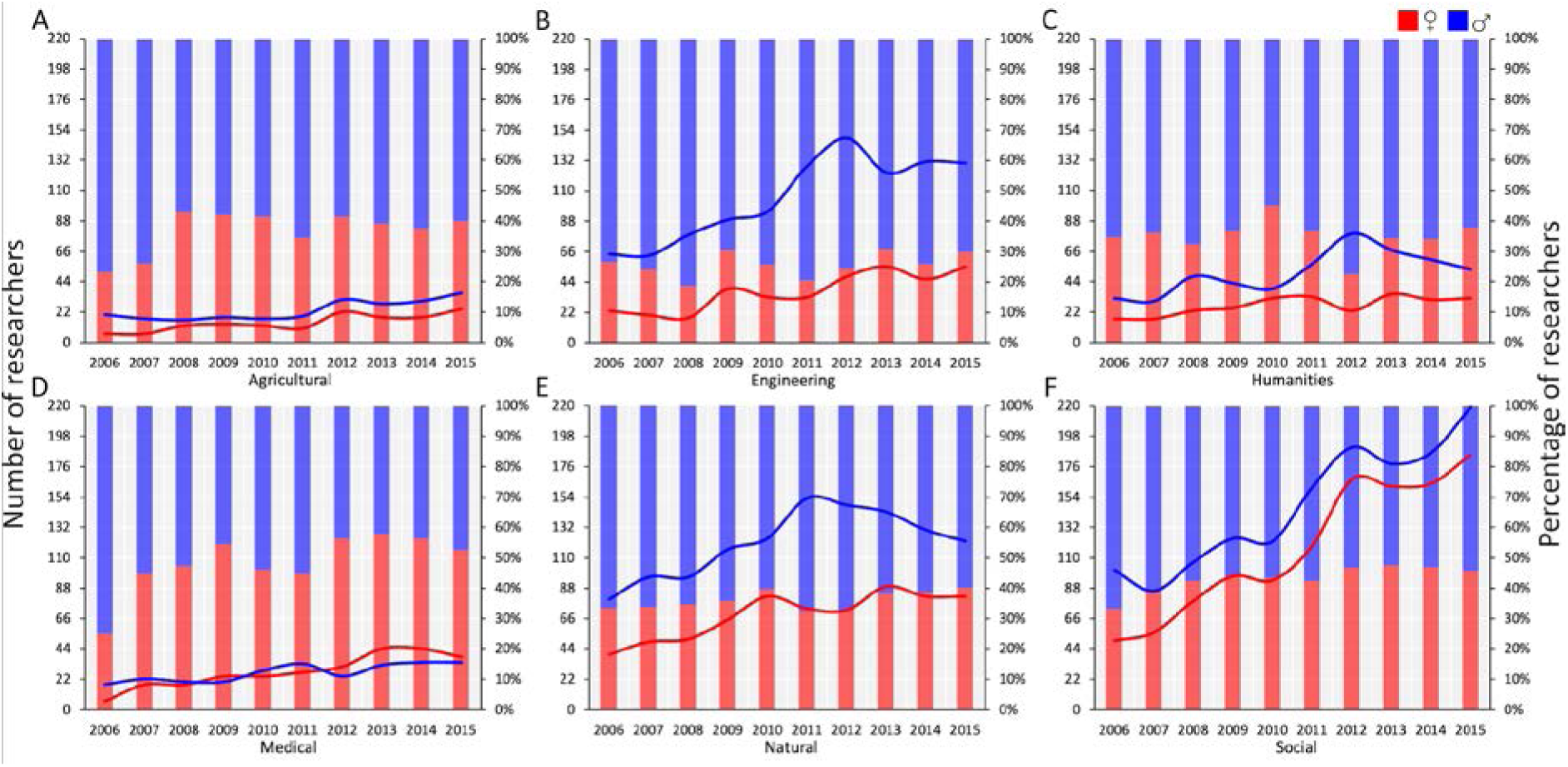
Female (red) and male (blue) representation in doctoral graduates in Colombia between 2000 and 2015, represented as number of researchers (lines) and percentage of total researchers (bars). Results are divided into six different research fields: Agricultural (A), engineering (B), humanities (C), medical (D), natural (E), and social (F).

Individual linear regressions showed no significant correlation between the increase in female doctoral graduates and the overall percentage of female researchers across research areas (Table 1). From 2006 to 2015, access to scholarships for postgraduate studies showed higher women representation for master’s degrees (49.10%, Fig. 7A) than for doctoral degrees (40.46%, Fig. 7B). However, women representation in scholarships for doctoral studies had the highest increase (4.06%), compared to master’s degrees (3.30%).

**Table 1.**
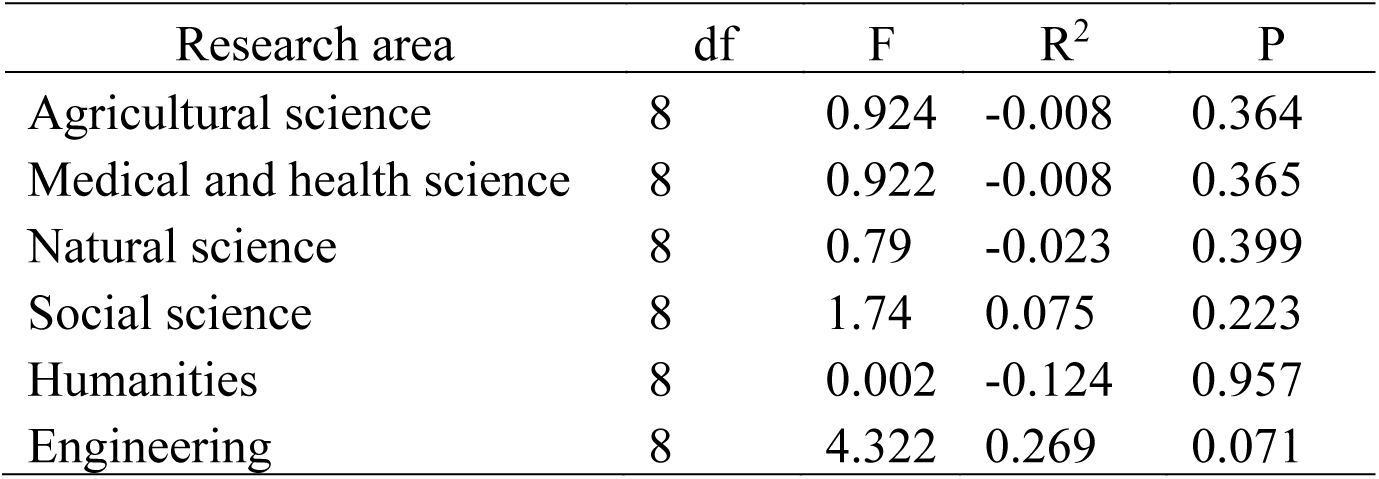
Results of linear regression modelling analysis of the interaction between female doctoral graduates and overall percentage of women in six different research areas.

**Figure 7.**
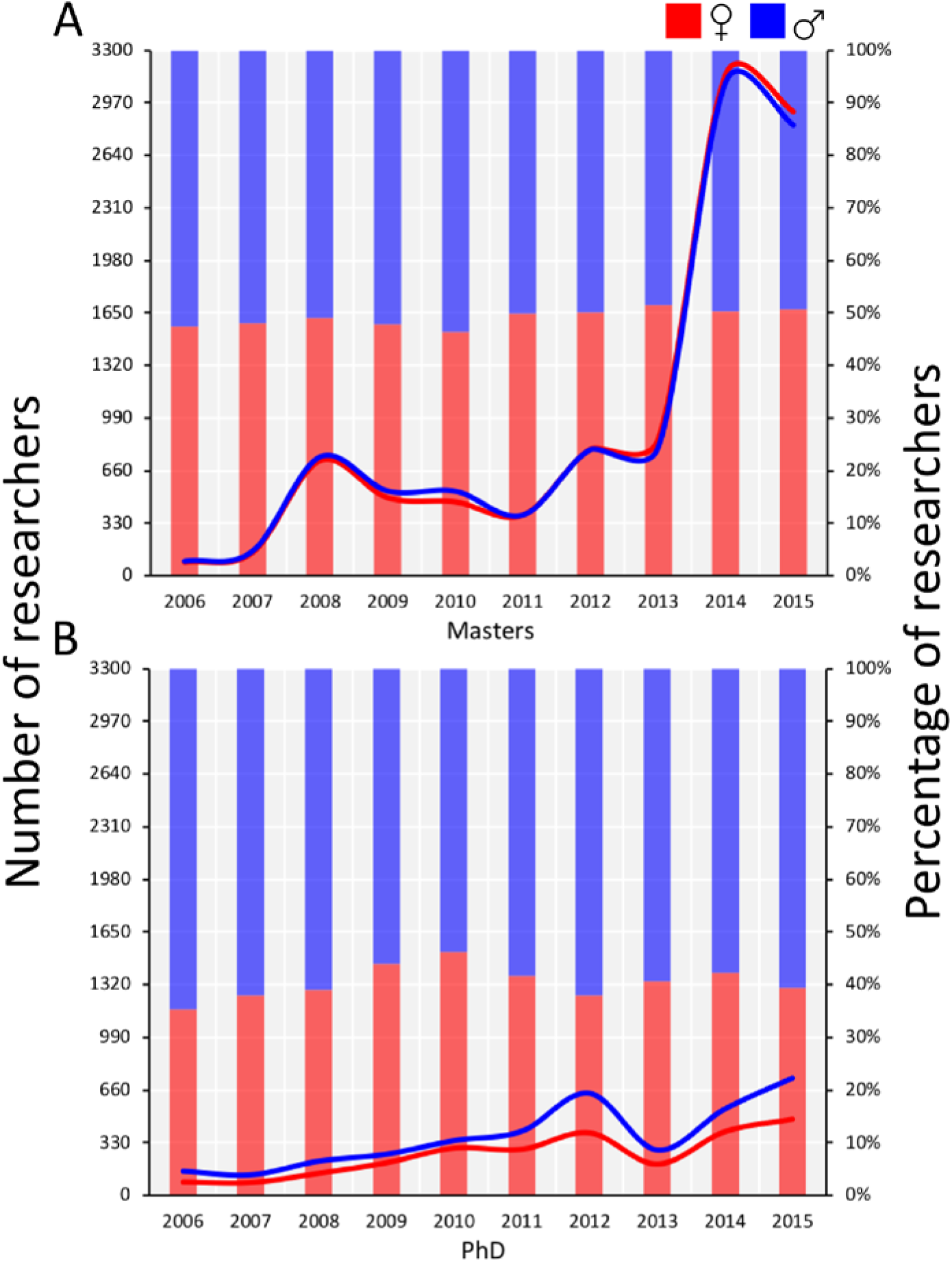
Women (red) and men (blue) access to scholarships for postgraduate studies in Colombia between 2000 and 2015, represented as number of scholarship awardees (lines) and percentage of total awardees (bars). Scholarships for masters’ degrees (A) and doctoral studies (B).

For the 2000 to 2015 period, a positive and statistically significant correlation was found between the percentage of women in science and the percentage of GDP invested in R&D (*p* < 0.001, *r*^2^ = 0.769). Based on logistic function modelling, the projections of future women representation in science predict that gender parity in the science workforce in Colombia could take up to 50 years (Fig. 8A). Decomposed based on research field, projections indicate that gender parity can take from three years (humanities) to more than 200 years (engineering). Medical sciences represent the only scenario were projections indicate a decrease in women representation. Years until gender parity in access to scholarships for postgraduate studies could range from two (social and agricultural) to 50 years (engineering), with the exceptional case of the humanities, where women representation is predicted to decrease (Fig. 8B). Finally, temporal projections to gender parity across researcher rank levels suggested that the junior rank could be first to reach gender parity, followed by emeritus, associate and senior ranks, where estimated years to parity range from five to 90 years (Fig. 8C). Raw data is available as a supplement.

**Figure 8.**
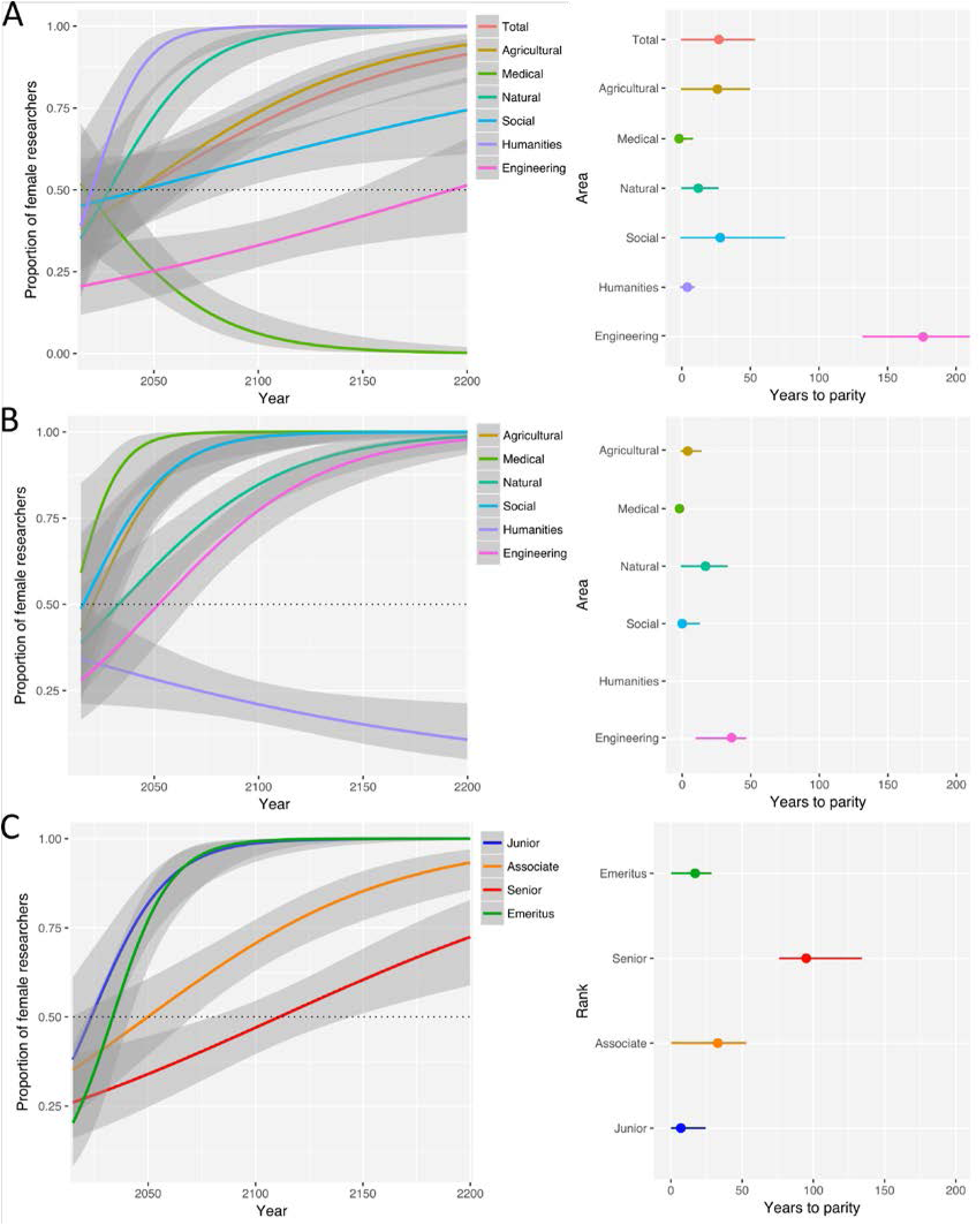
Temporal projections with 95% confidence intervals of women representation in science between 2000 and 2200 based on logistic function modelling using a binomial distribution, as a proportion of total researchers. Projections are represented as temporal trajectories of women representation (left column), and as the amount of years to parity from 2018 (right column). Projections were made for the overall science workforce (A) and access to scholarships for postgraduate studies (B) for six different research fields, and overall science workforce across researcher rank levels (C).

## Discussion

This study represents the more comprehensive study of the status of gender parity in science in Colombia in the 21st century (Daza & Bustos, 2008), providing a diagnosis of the recent trends of women representation across research fields, researcher ranks and education level, with some estimates of future temporal projections to gender parity. The analyses performed in this study represent to the best of my knowledge the first quantitative study examining the official data gathered by COLCIENCIAS and made freely available through the SCIENTI platform starting from 2015 (COLCIENCIAS, 2017). This study also builds on previous results that have helped elucidate the reality of gender pay gap (Franco-Orozco & Franco-Orozco, 2018), access to education and scholarships in Colombia. Given the scarcity of similar studies examining gender parity in science for the Latin America and the Caribbean region, the current study also helps to inform the regional context of women in science, associating it with other recent studies in Colombia and Brazil (Franco-Orozco & Franco-Orozco, 2018; Valentova et al., 2017).

The results presented here show widespread lack of gender parity and underrepresentation of women in science in Colombia across the 21st century, informing previous analyses that reported an extensive gender pay gap in science-related work fields (Franco-Orozco & Franco-Orozco, 2018). Following similar trends found for other countries, the lowest level of women representation was found in engineering, an area heavily dominated by implicit gender stereotypes, followed by the humanities and natural sciences (Ceci & Williams, 2011; Ceci et al., 2009; Christie et al., 2017; Franco-Orozco & Franco-Orozco, 2018; Meyer et al., 2015; Valentova et al., 2017). Gender inequality in the humanities has been reported in salaries and tenure promotion in the US, showing a lack of correlation with productivity and pointing to subconscious gender bias as a possible influencing factor. However, the study of gender inequality in the humanities remains scarce, highlighting the need for increased research efforts. Recently, gender disparity in the natural sciences has been increasingly studied, providing evidence of gender differences in the length and tone of recommendation letters (Dutt et al., 2016), participation at scientific events (Débarre et al., 2018; Jones et al., 2014), and representation in editorial boards (Cho et al., 2014). In Colombia, previous studies have provided evidence of a gender pay gap in the areas aforementioned, both for undergraduate and postgraduate graduates (Franco-Orozco & Franco-Orozco, 2018). Temporal trends in the gender gap in Colombia followed similar trends found in other countries (Christie et al., 2017; Ramakrishnan et al., 2014; Valentova et al., 2017; van den Besselaar & Sandstrom, 2017; van der Lee & Ellemers, 2015). The decrease in women representation in medical and health science contrasts with the fact that it was the only research area to have reached gender parity in Colombia. Despite a general level of gender parity across different countries (Franco-Orozco & Franco-Orozco, 2018; Ramakrishnan et al., 2014; van den Besselaar & Sandstrom, 2017), women underrepresentation in the medical and health science could still be found at more senior positions, illustrating the ‘leaky pipeline phenomenon’ (Ramakrishnan et al., 2014). A decreasing representation of women in medical sciences, generally considered a gender equal field, highlights the need to implement initiatives that not only promote the participation of women in male-dominated research areas, but also secure the retention of women as they move upward to more senior rank levels.

Since it is expected that the average age of researchers increases at higher researcher rank levels, the lower average age in female researchers could reflect the lower representation of women in the most senior research levels, signalling another potential impact of the ‘leaky pipeline phenomenon’ (Blickenstaff, 2005; Pell, 1996). However, lower age in women could also represent an opportunity to secure the retention of a younger population of female researchers, driving a future increase in the representation of women at more senior levels as they move upward across research ranks. Moreover, the underrepresentation of women as a proportion of scholarship awardees for doctoral studies and doctoral graduates indicate that the completion of doctoral studies might be a limiting factor influencing the loss of women beyond the junior researcher rank (Franco-Orozco & Franco-Orozco, 2018; Valentova et al., 2017; van den Besselaar & Sandstrom, 2017). Also, the lack of a significant correlation between female doctoral graduates and the percentage of female researchers indicate that retaining doctoral graduates is a key component to consider. Based on this, it can be hypothesised that a boost in the proportion of women with doctoral degrees in a younger population of female researchers could have a cascading effect, encouraging the participation and retention of women across ranks (Shen, 2013).

Based on data from the UIS, Colombia ranks 15th out of 20 Latin American countries with available data, sitting below the average of women participation in Latin America (45%), distant from countries like Bolivia (63%), Venezuela (62%) and Trinidad and Tobago (54%), countries with the highest percentage of women representation in science in the region (UNESCO Institute of Statistics, 2018). Considering the state of political and civil unrest that has prevailed over the last decades, it could be argued that the long-lasting internal armed conflict could be one of the main drivers influencing women participation in Colombian science (Daza & Bustos, 2008; Franco-Orozco & Franco-Orozco, 2018). It is estimated that 3.5 million women were victims of the internal conflict (49.5% of the victims), and that between 2010-2015, more than 800,000 were victims of some kind of sexual violence (Cifelli & Diaz, 1989; International, 2017; Pérez, 2008). Moreover, data from 2000 from the United Nations Development Program estimated that between 60-70% of Colombian women have suffered some kind of violence. According to the 2018 Global Peace Index report (ranking the intensity of the internal conflict of a country), Colombia ranks 145^th^ of 163 countries studied (Institute for Economics and Peace, 2018). Nonetheless, comparing the percentage of women in science in Colombia with other countries with similar intensity of internal conflict, Colombia ranks 9^th^ in 20 countries (Table 2), five points above the average for these countries (32.65%). This could indicate that despite the differential impact that war has on women’s rights, internal conflict is not the only limiting factor leading to women underrepresentation in science, and so additional factors should also be considered.

**Table 2.**
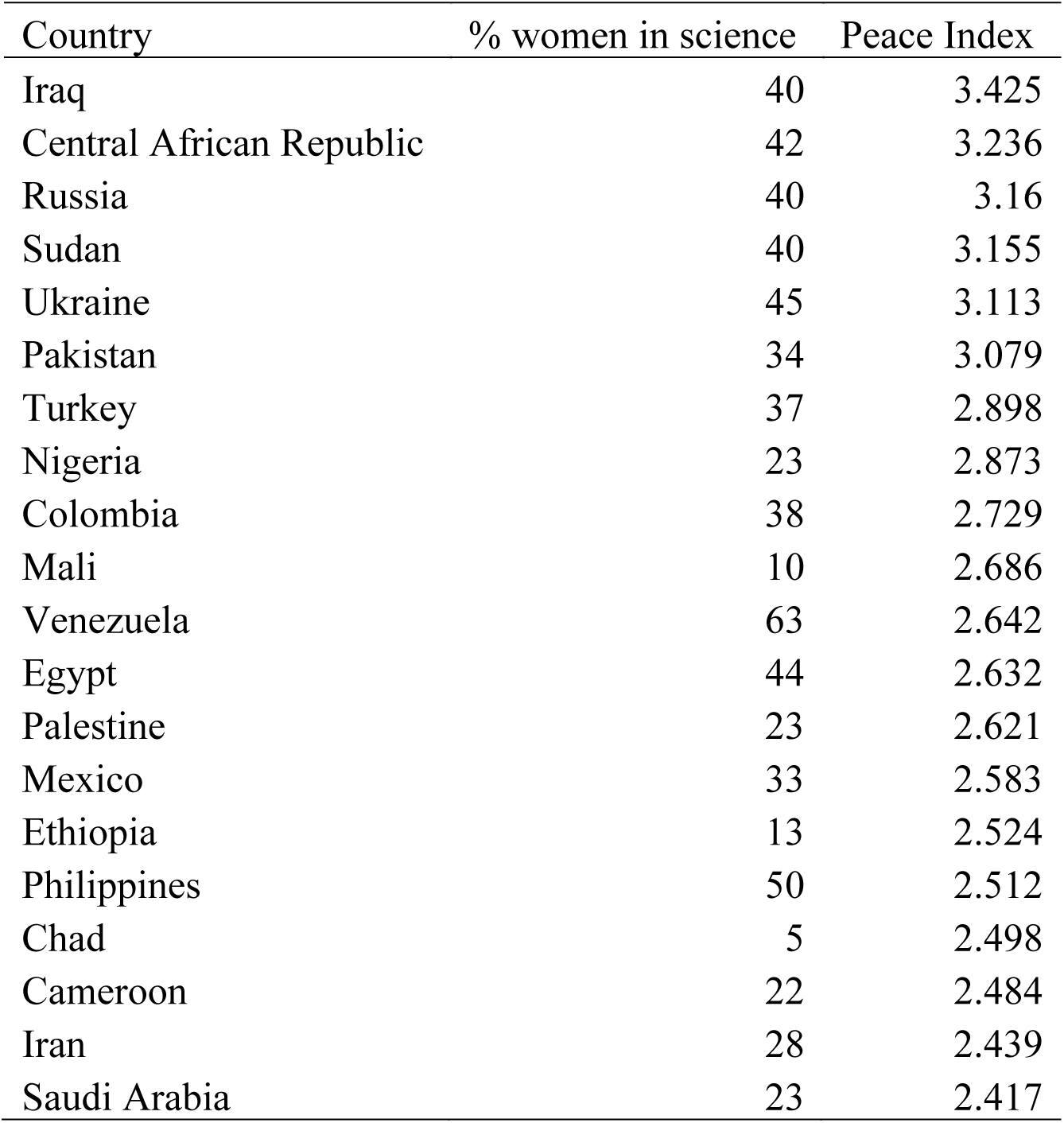
Comparison of percentage of women participation in science between 20 countries with similar 2018 Global Peace Index. Country % women in science Peace Index

R&D expenditure has been discussed as a potential driver of women underrepresentation in science (Ceci & Williams, 2011; Christie et al., 2017; van den Besselaar & Sandstrom, 2017). Despite an increase in the percentage of GDP invested in R&D between 2000 (0.13) and 2013 (0.27) (World Bank, 2018), COLCIENCIAS has seen a steady decrease in its annual budget between 2013 (430,000 million COP) and 2018 (337,000 million COP) (COLCIENCIAS, 2017). The present results did show a significant correlation between women representation and R&D expenditure between 2000-2015, suggesting that the decrease in the annual budget of COLCIENCIAS since 2013 could have played a part in the decrease in the percentage of women in science in Colombia between 2012 and 2015. R&D expenditure in Colombia is below the average for Latin America and the Caribbean (0.7%) and is the lowest of the five countries with the highest scientific output in the region (UNESCO Institute of Statistics, 2018), which reinforces the need for an increase in R&D expenditure to tackle women underrepresentation in the future. Moreover, comparing the percentage of women in science in countries with similar R&D expenditure (0.21 to 0.36% of GDP), the percentage of women in science in Colombia is below average (39.69%, Table 3). This informs the results above and suggests that the internal conflict and lack of funding are not the only decisive factors that could explain gender disparity in science in Colombia.

**Table 3.** Comparison of percentage of women participation in science between countries with percentage of GDP destined to R&D between 0.21 and 0.36, based on data from World Bank (2018).

| Country | % of women in science | R&D expenditure |
| --- | --- | --- |
| Uruguay | 50 | 0.36 |
| Mozambique | 29 | 0.34 |
| Georgia | 52 | 0.32 |
| Mali | 10 | 0.31 |
| Colombia | 38 | 0.29 |
| Eswatini | 41 | 0.27 |
| Armenia | 52 | 0.25 |
| Oman | 28 | 0.25 |
| Pakistan | 34 | 0.25 |
| Venezuela | 63 | 0.25 |
| Bosnia and Herzegovina | 47 | 0.22 |
| Bermuda | 32 | 0.22 |
| Uzbekistan | 40 | 0.21 |

The legal framework ruling the institutional procedures that promote gender parity in science is also a major mechanism for the enhancement of women representation (Ceci & Williams, 2011; Ceci et al., 2009; Pell, 1996). Recently, Colombia has made major steps towards ensuring the protection of women’s rights, especially in the context of the internal armed conflict (Overseas Development Institute, 2015). Law 581 of 2000 established a minimum quota of 30% of women representation in government (Bustamante, 2007). A revised quota law was established in 2011 (law 1475 of 2011), extending the implementation of the 30% quota to the formation and operation of political parties and political movements (Bustamante, 2007). However, the impact of the measurable benefits derived from these laws is under debate (Batlle, 2016). A battery of additional laws has been established over the last decade, protecting the principle of gender equality, access to land and access to justice in cases of sexual violence. Nevertheless, despite the advancement in the protection of gender parity and women’s rights, no legislation has been established governing the representation of women in science. As an attempt to promote, stimulate and highlight women participation science and technology in Colombia, the Colombian Network of Scientific Women (Red Colombiana de Mujeres Científicas) was created in 2015 (RCMC, 2015). Examples of legislation promoting gender parity in science can be drawn from countries such as Spain (Law 14 of 2011), the United States of America (the Promoting Women in Entrepreneurship Act and the INSPIRE Women Act), and the European Union (article 16 of the Regulation 1291 of 2013 ruling the Horizon2020 program). Current discussions of science legislation in Colombia have focused on a proposal for the creation of a Ministry of Science, Technology and Innovation, by changing Law 1286 of 2009 (El Espectador, 2018a). This would elevate COLCIENCIAS from an administrative department under the National Planning Department to an independent ministry. Beyond the discussion around the status of the institution, COLCIENCIAS has currently been under public scrutiny mainly due to the instability in its leadership, evidenced by the 10 directors that have been named since 2010 (El Espectador, 2018b).

Assessing and controlling for unconscious bias is also crucial to diminish gender disparity in academia, as it addresses the cultural and psychological drivers of women underrepresentation (Ceci & Williams, 2011; Christie et al., 2017). Extensive evidence currently available on the sources of unconscious bias that impact the participation of women in science could be divided into two components: opposite gender exclusion and self-exclusion (Ceci & Williams, 2011; Christie et al., 2017). Opposite gender exclusion could be described as the result of a tendency to favour people of the same gender, leading to the unintentional exclusion of the opposite gender (Murray et al., 2018). This phenomenon has been reported equally for women and men in science (Murray et al., 2018). However, given the heavily male-dominated demographics of the STEMM workplace, women underrepresentation in science could be partly due to an exacerbated opposite gender exclusion. Implementing double-blind peer review and increasing gender and international diversity in review committees have been proposed to enhance the representation of minorities both in scientific publishing and in more senior institutional positions (Murray et al., 2018). Self-exclusion in science, on the other hand, can be viewed as the tendency to restrict oneself ‘s gender from involving in scientific activities, resulting from a variety of unconscious negative stereotypes (Moss-Racusin et al., 2012; Smeding, 2012).

Recent research has showed that negative gender stereotypes on intellectual prowess appear early during childhood, leading both boys and girls to consider men as more intelligent than women by age six (Bian et al., 2017). This early predisposition does not reflect a natural tendency in any way, as recent findings have found higher average academic grades for girls than boys (O’Dea et al., 2018). The same study estimated gender parity in the top 10% of a STEMM-related class, and higher women representation in non-STEMM-related classes (O’Dea et al., 2018). Self-exclusion has also been reported in later stages of the academic career, in faculty members of different Science faculties at University level (Moss-Racusin et al., 2012). Both female and male faculty members showed a tendency to rate male students higher than female students, favouring higher salaries and mentoring for male applicants, leading to a lesser probability for female students to be hired (Moss-Racusin et al., 2012).

Additionally, pre-existing bias was associated with less support for female students but did not associate with reactions to male students (Moss-Racusin et al., 2012). This suggests that unconscious bias is a major driver of women’s exclusion in science, both as a result of opposite gender exclusion and self-exclusion.

## Conclusions

The results presented here elucidate the state of women participation in science across the 21^st^ century, highlighting a generalised trend of women underrepresentation. Even though temporal trends show an increase in the percentage of women across all but one research area (i.e. medical and health science), greater efforts are needed to increase and retain gender parity across research fields. Initiatives to retain women in Colombian science should be of special focus for the medical and health science, as it is the only research area to both have reached gender parity and show a steady decrease in women representation since 2010. Given the lower percentage of female researchers in engineering, humanities and natural sciences, this should be areas of special focus for institutions in research and education. The lower average age of female researchers could represent an opportunity to address the ‘leaky-pipeline phenomenon’, ensuring that young female researchers are supported as they move upward to more senior research levels. Our results suggest that improving access to scholarships for doctoral studies, and the retention of female doctoral graduates in research, could be major strategies to ensure the increase women representation in science in the future. Nonetheless, equal efforts should be made to improve the career prospects and working environment of Colombian women scientists in the present. This study also highlights the importance of long-term monitoring of demographic trends in science, in order to inform individual, institutional, governmental and global initiatives focused on increasing gender parity in STEMM. Following the increasing understanding of discrimination in science (Hughes, 2018; Pew Research Centre, 2018), future studies and discussion should also expand to evaluate representation of racial, ethnic and sexual minorities to inform the prevalence of minority discrimination in Colombian science. More and more refined data would allow more robust modelling techniques to be implemented in the estimates of temporal projections for gender parity, improving our predictive power. Consequently, the temporal projections presented here should be taken with caution as they are only a statistical representation of the data available. Without a greater overarching commitment to monitor and strengthen women representation in STEMM at a regional scale, not only in Colombia but globally, gender disparity could remain a critical issue that will plague science for decades and even centuries to come.

## Supporting information

## Acknowledgements

This work was inspired by countless Colombian female scientists who have endured the difficulties of an armed conflict, not only to challenge gender stereotypes and inspire other women to pursuit STEMM-related careers, but also to encourage male scientists to evaluate and reconsider our role in society and to defy our own stereotypes. Special thanks to Laura Castañeda-Gómez and Ana López-Aguirre for improving several early versions of the manuscript.

